# WASP: allele-specific software for robust discovery of molecular quantitative trait loci

**DOI:** 10.1101/011221

**Authors:** Bryce van de Geijn, Graham McVicker, Yoav Gilad, Jonathan K. Pritchard

**Affiliations:** Department of Human Genetics, University of Chicago; Committee on Genetics, Genomics and Systems Biology, University of Chicago; Department of Genetics, Stanford University; Department of Biology, Stanford University; Howard Hughes Medical Institute, Stanford University

## Abstract

Allele-specific sequencing reads provide a powerful signal for identifying molecular quantitative trait loci (QTLs), however they are challenging to analyze and prone to technical artefacts. Here we describe WASP, a suite of tools for unbiased allele-specific read mapping and discovery of molecular QTLs. Using simulated reads, RNA-seq reads and ChIP-seq reads, we demonstrate that our approach has a low error rate and is far more powerful than existing QTL mapping approaches.

---

Next generation sequencing data can be used to identify allele-specific signals because reads that overlap heterozygous sites can be assigned to one chromosome or the other. Molecular QTLs are associated with allelic imbalance^1–4^, and thus allele-specific reads can potentially augment the power of statistical tests for QTL discovery^5^. However, use of allele-specific reads can introduce artefacts into many stages of analysis. Uncorrected mapping of allele-specific reads can be highly biased and can easily yield false signals of allelic imbalance^6,7^. Homozygous sites which are incorrectly called as heterozygous are another source of false positives, and allele-specific read counts are overdispersed compared to the theoretical expectation of a binomial distribution^8^. Here we describe a suite of tools called WASP that is designed to overcome these technical hurdles. WASP carefully maps allele-specific reads, corrects for incorrect heterozygous genotypes and other sources of bias, and models overdispersion of sequencing reads. Finally, by integrating allele-specific information into a QTL mapping framework WASP attains greater power than standard QTL mapping approaches.

**Figure 1.**
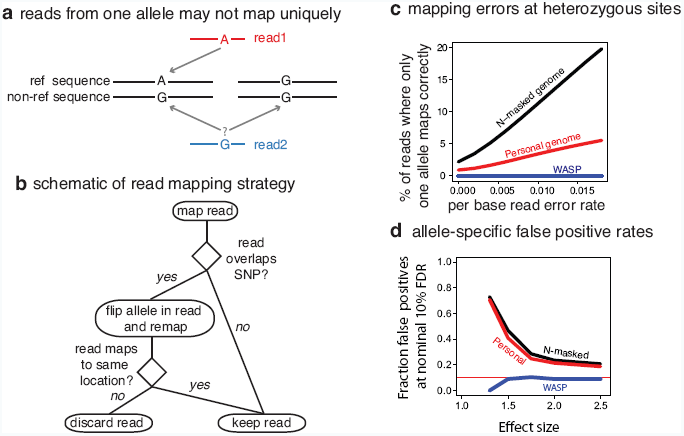
Mapping of allele specific reads. (**a**) Mapping to ‘personalized’ genomes can result in allelic bias because reads from one allele may not map uniquely. (**b**) Schematic of mapping pipeline to remove allelic bias. (**c**) The percentage of simulated 100 bp reads at heterozygous sites where a read with one allele maps correctly and the corresponding read with the other allele does not. Reads were simulated with sequencing errors introduced at several different rates. (**d**) The fraction of false-positives as a function of the effect size using a nominal Benjamini-Hochberg false-discovery rate of 10%. We simulated 100 bp allele-specific reads under null (odds-ratio = 1) and alternative models (odds-ratio > 1) of allelic imbalance at heterozygous sites in the genome. 90% and 10% of sites were assumed to be null and alternative sites respectively. We mapped reads using WASP or the personal- or N-masked mapping strategies and called allele-specific sites using a binomial test.

Mapping of reads to a reference genome is biased by sequence polymorphisms^6^. Reads which contain the non-reference allele may fail to map uniquely or map to a different (incorrect) location in the genome^6^. A common approach is to map to a ‘personalized’ genome where the reference sequence is replaced by non-reference alleles that are known to be present in the sample^9^. However, personalized genomes do not fully address the mapping problem because the genomic locations that are uniquely mappable in the reference and non-reference genome sequences differ (Fig. 1). While these type of errors may only affect a small number of sites, they comprise a large fraction of the most significant results when tests of allelic imbalance are performed genome-wide (Fig. 1).

WASP uses a simple approach to overcome mapping bias that can be readily incorporated into any read mapping pipeline. First, reads are mapped normally using a mapping tool selected by the user; mapped reads that overlap single nucleotide polymorphisms (SNPs) are then identified. For each read that overlaps a SNP, its genotype is swapped with that of the other allele and it is re-mapped. If a re-mapped read fails to map to exactly the same location, it is discarded (Fig. 1). Unknown polymorphisms in the sample are not considered but will typically have little effect since the tests of allelic imbalance are only performed at known heterozygous sites.

We evaluated the performance of WASP’s remapping method by simulating reads at heterozygous sites in a lymphoblastoid cell line that has been completely genotyped and phased (GM12878). At each heterozygous SNP we simulated all possible overlapping reads from both haplotypes, additionally allowing reads to contain mismatches at a predefined sequencing error rate. We mapped the simulated reads using three approaches to account for mapping bias: mapping to a genome with N-masked SNPs, mapping to a personalized genome, and mapping to the genome using WASP. While reads mapped to the N-masked and personalized genomes were substantially biased and resulted in a large number of false positives, reads mapped using WASP were almost perfectly balanced (Fig. 1).

WASP employs a number of techniques to remove noise and biases from mapped reads. Amplification bias is a common feature of experiments that yield libraries with low complexity (e.g. ChIP-seq). To control for amplification bias it is common to remove ‘duplicate’ reads that map to the same location. However, existing tools that remove duplicate reads retain the one with the highest mapping score, which will usually match the reference^10^. WASP provides a tool to filter duplicate reads at random, thus eliminating reference bias from this step.

GC content often affects read depth in a manner that is inconsistent between sequencing experiments^1,11^. In addition, the distribution of read depths across the genome differs from experiment to experiment. For example, ChIP-seq experiments with more efficient pull-downs tend to have more reads within peaks. WASP corrects for both of these issues by fitting polynomials to the genome-wide read counts and calculating a corrected read depth for each region (see Methods).

Both allele-specific and total read depth counts are more dispersed than expected under models of binomial and poisson sampling^8,12^. To accommodate overdispersion in the data, WASP estimates separate overdispersion parameters for each individual and genomic region used in a study. Finally, to account for any remaining unknown covariates, WASP allows principal components to be included in the model fitting procedure (see Methods).

**Figure 2.**
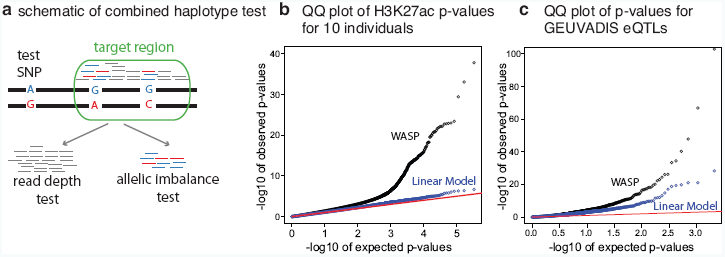
The combined haplotype test and its performance. (**a**) Schematic of the combined haplotype test. A ‘test SNP’ is tested for association with mapped reads within a ‘target region’. All reads are used by the read depth component of the test; allele-specific reads are used by the allelic imbalance component of the test. (**b**) Identification of novel QTLs using H3K27ac ChIP-seq data from 10 Yoruba lymphoblastoid cell lines. (**c**) Identifying European eQTLs from the GEUVADIS consortium using an independent dataset of RNA-seq from 69 Yoruba lymphoblastoid cell lines.

Following correction for biases described above, WASP uses a statistical test, the combined haplotype test (CHT), to identify cis-acting QTLs. The CHT tests whether the genotype of a ‘test SNP’ is associated with total read depth and allelic imbalance in a ‘target region’ (Fig. 2). The CHT jointly models two components: the allelic imbalance at phased heterozygous SNPs and the total read depth in the target region. The two components of the test are linked together by shared parameters that define their effect sizes.

For a target region and test SNP pair, the CHT models the expected number of reads for an individual as a function of the individual’s genotype, the effect size, the GC content, additional covariates (such as principal component loadings), and the total number of mapped reads in the region (across all individuals). The probability of the observed number of reads in the target region is calculated using the expected number of reads and two overdispersion parameters.

Allelic imbalance of reads overlapping heterozygous SNPs within a target region is modeled as a function of the shared effect size parameters. The probability of the observed allelespecific read counts is then defined by the effect size and a single overdispersion parameter. We also allow for the possibility of genotyping errors by assuming that allele-specific read counts are drawn from a mixture, with a small probability that a given individual is a mistyped homozygote.

To evaluate the performance of WASP on a small dataset, we used it to call novel QTLs genome-wide using data from H3K27ac ChIP-seq experiments that were performed in 10 lymphoblastoid cell lines^12^. Remarkably, WASP identifies 2426 H3K27ac QTLs (10% FDR), whereas a linear regression approach is unable to identify any (Fig. 2).

We also evaluated the ability of WASP to call gene expression QTLs (eQTLs) in a larger dataset. We obtained a set of 2098 expression QTLs (eQTLs) identified in 462 lymphoblastoid cell lines (LCLs) derived from European individuals^13^. We tested whether we could identify these eQTLs, using an independent dataset of RNA-seq from 69 Yoruba LCLs^1^. WASP discovers 627 of these eQTLs at a false discovery rate (FDR) of 10%, which is impressive considering (1) our smaller sample size, (2) that some fraction of the original eQTLs are false positives, and (3) that some of the European eQTLs will be absent or at very low frequency in the Yoruba (Fig. 2). This number increases to 673 when 5 principal components are included as covariates. By comparison, when we adopt a standard eQTL discovery method (linear regression on quantile normalized and GC-corrected data), we identify only 446 eQTLs (617 when 5 principal components are included as co-variates). *P*-values obtained by running the CHT on the same dataset with permuted genotypes do not depart substantially from the null expectation, indicating that the test is well-calibrated (Supplementary Fig. S4).

These results demonstrate that WASP is a powerful approach for the identification of molecular QTLs, particularly when sample sizes are small. WASP accounts for numerous biases in allele-specific data and is flexible enough to work with different read mappers and multiple types of sequencing data such as ChIP-seq and RNA-seq. By modeling biases and dispersion differences directly, WASP eliminates the need for quantile normalization of the data, thereby making estimated effect sizes more interpretable.

The source code and documentation for WASP are open source and can be downloaded from https://github.com/bmvdgeijn/WASP/.

## Acknowledgements

This work was supported by the Howard Hughes Medical Institute, NIH grants HG007036, HG006123, MH101825, and GM007197 and by a NSF Graduate Research Fellowship (DGE-0638477) to BvdG. We would like to thank X. Shirley Liu’s lab for hosting GM as a visitor in the Department of Biostatistics and Computational Biology at the Dana-Farber Cancer Institute while this work was conducted. We thank members of the Liu, Pritchard, Stephens and Gilad labs for helpful discussions.

